## Supplementary material for "Stiffness and tension gradients of the hair cell’s tip-link complex in the mammalian cochlea"

The following figure and table supplements are available for figure 2:

**Figure supplement 1.** Velocity field of a fluid jet.

**Figure supplement 2.** Geometrical characteristics of a fluid jet.

**Figure supplement 3.** Rise time and linearity of the fluid-jet stimulus.

**Figure supplement 4.** Test of fluid-jet calibration in the frog's sacculus.

**Figure supplement 5.** Mechanical creep during a force step.

**Table supplement 1.** Statistical significance.

The following figure and table supplements are available for figure 3:

**Figure supplement 1.** Gating-spring contribution to the hair-bundle stiffness.

**Figure supplement 2.** Hair-bundle morphology along the tonotopic axis.

**Figure supplement 3.** Transduction currents and number of intact tip links along the tonotopic axis.

**Table supplement 1.** Morphological parameters of inner and outer hair-cell bundles.

**Table supplement 2.** Statistical significance.

The following table supplement is available for figure 5:

**Table supplement 1.** Statistical significance.

The following table supplement is available for figure 6:

**Table supplement 1.** Statistical significance.

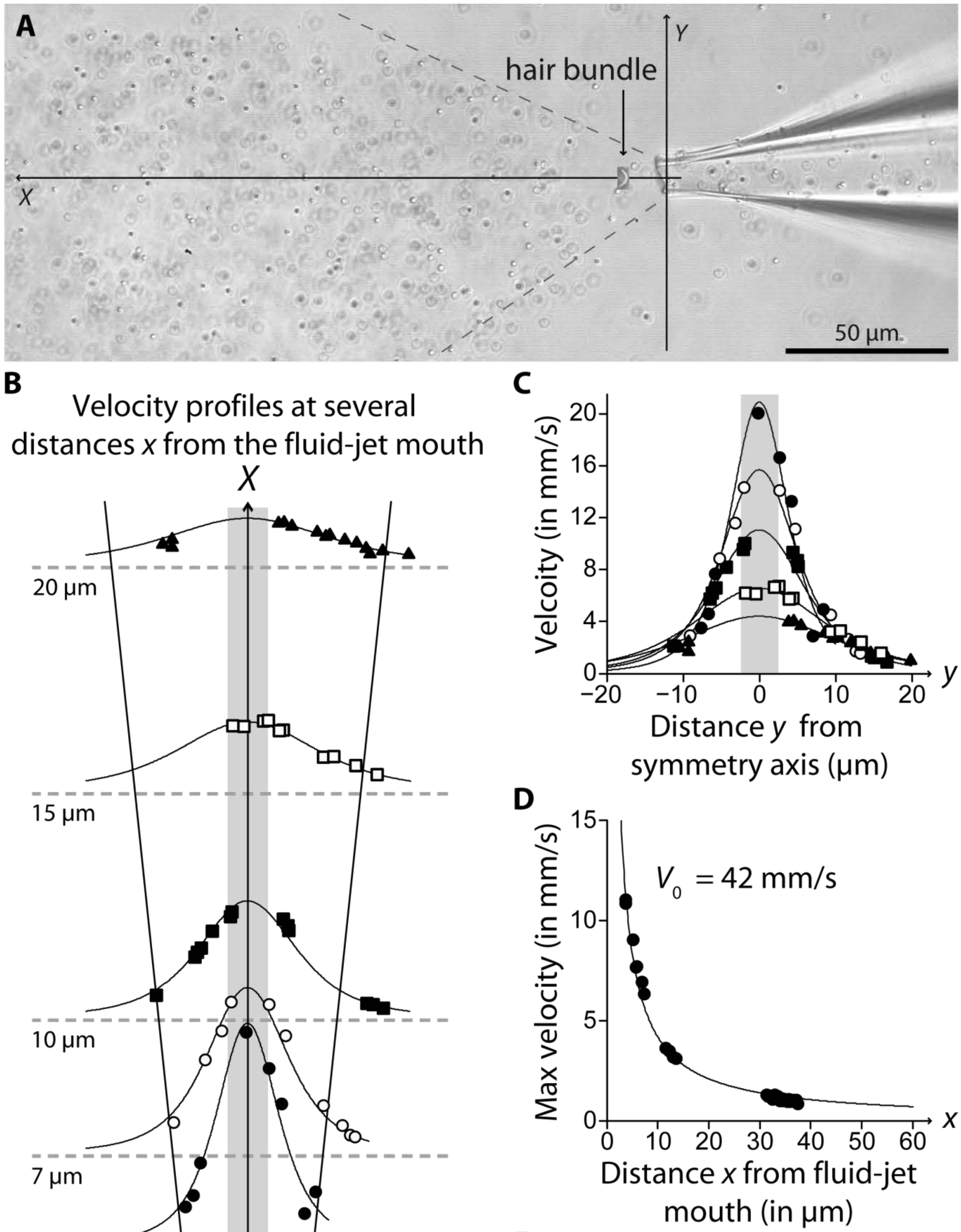


E

| Distance from pipette (µm) | $V_{max}$ (mm/s) | $A$ (µm) | $\overline{\alpha}$ (°) |
|---|---|---|---|
| ● 5 | 21 | 6 | 32 |
| ○ 7 | 16 | 9 | 31 |
| ■ 10 | 11 | 11 | 31 |
| □ 15 | 7 | 16 | 36 |
| ▲ 20 | 4 | 19 | 36 |

Hair bundle width

$\alpha$ = 30° cone

**Figure 2–figure supplement 1: Velocity field of a fluid jet.** **(A)** Micrograph showing 200-nm beads entrained by a fluid jet; the beads were used as tracers for velocimetry. The dotted lines delimit the fluid cone coming out of the pipette; its half-aperture $\alpha = 30°$ was in agreement with that measured with Coomassie blue (Figure 2–figure supplement 2), considering that the diameter of the fluid-jet pipette was here 10 μm. A scaled picture of an outer hair-cell bundle was inserted in the micrograph to illustrate how a hair bundle was positioned within the fluid jet. A movie was recorded with a high-speed camera (Photron Fastcam Mini UX50) at 5,000 images/s. The position of the beads was automatically tracked in the $XY$ plane using the TrackMate plugin (Tinevez et al., 2017) of the image-processing software Image J (National Institute of Health, Bethesda, USA). **(B)** Longitudinal-velocity profile $v_X(x, y)$ along the transverse $Y$ axis, at several values of the distance $x$ from the pipette mouth along the $X$ axis. The data were well described by the function $v_X(x, y) = V_{\mathrm{max}}(x)/(1 + (y/A(x))^2)^2$ (solid lines). The shaded area indicates the width of a hair bundle. We noticed that the half-aperture $\alpha$ of the fluid jet that we measured with Coomassie blue (Figure 2–figure supplement 2) or with the beads **(A)** was well approximated by $\alpha \cong \bar{\alpha} = \tan^{-1}\big((A(x) - D_{\mathrm{FJ}}/2)/x\big)$, where $D_{\mathrm{FJ}}$ is the diameter of the pipette; the oblique lines shown in the figure delimit the fluid jet and match the cone shown in **A**. As a result, when a calibration fiber was placed perpendicular to the fluid jet (Figure 2–figure supplement 2B), the fiber experienced viscous drag over a typical length $L(x) \cong 2A(x)$. **(C)** The same velocity profiles as those shown in **B** are here overlapped; different symbols correspond to beads at different distances $x$ from the mouth of the fluid-jet pipette, according to the legend shown in **B**. **(D)** Fluid velocity at $y = 0$ as a function of the distance $x$. The data was well described by $v_X(x, 0) = V_0/x$, with $V_0 = 42$ mm/s (solid line). **(E)** Fit parameters $V_{\mathrm{max}}(x)$ and $A(x)$ for the fits shown in **B** and **C** as solid lines, as well as the approximation for the fluid-jet half aperture $\bar{\alpha}(x)$.

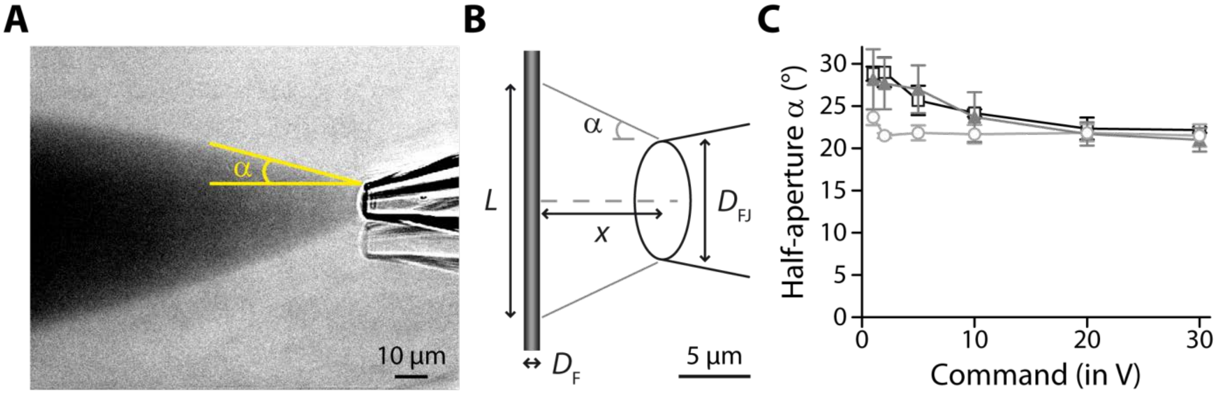


**Figure 2−figure supplement 2: Geometrical characteristics of a fluid jet.** **(A)** Visualization of a fluid jet using a solution containing a dye (Coomassie Brilliant Blue). To enhance contrast, a background image was recorded in the absence of the fluid-jet pipette and subtracted to the image of the fluid jet, resulting in the image shown here. The angle between the horizontal axis and the edge of the cone defines the half-aperture of the cone, noted α. Command voltage of −60 V. **(B)** Schematic representation of a calibration fiber intersecting a fluid jet over a length $\mathrm{L} = \mathrm{D_{FJ}} + 2x \tan \alpha$, in which $\mathrm{D_{FJ}}$ is the diameter of the pipette, $x$ is the distance from the pipette to the fibre and α is the half-aperture of the fluid jet, as defined in **A**. This length L and the diameter $\mathrm{D_F}$ of the cylindrical fiber is used to calculate the effective hydrodynamic radius of the fibre (see Methods in the main text). **(C)** Half-aperture α as a function of the command voltage to the fluid-jet device, for pipette diameters ranging between 3−5 µm (light gray circles; n = 3), 5−7 µm (gray triangles; n = 5) and 7−9 µm (black squares; n = 3). Error bars correspond to mean values ± SEM. In practice, the maximal command voltage that we used to probe hair-bundle stiffness was 4±0.3 V (mean±SEM; n = 139). Over this range, the half-aperture of the fluid jet was nearly constant. We took a value of 22° for pipettes with a diameter of 3−5 µm and of 27° for pipettes with a diameter of 5−9µm.

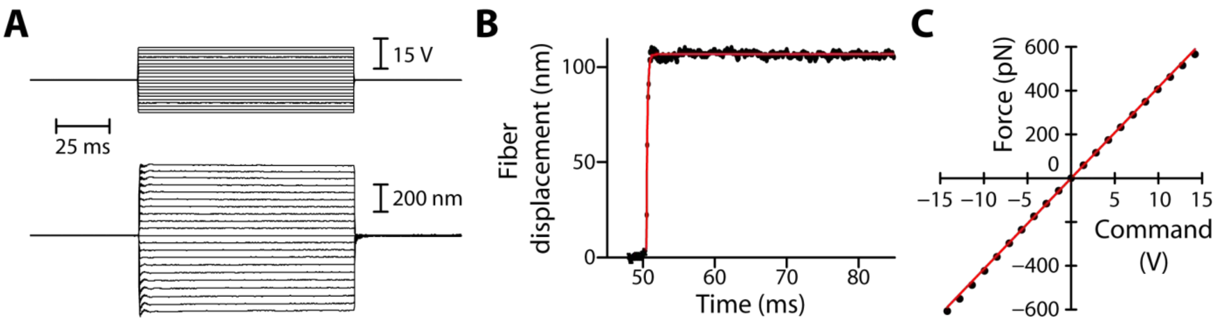


**Figure 2–figure supplement 3: Rise time and linearity of the fluid-jet stimulus. (A)** Deflection of a flexible fiber (bottom) in response to a series of command-voltage steps (top) applied to the fluid-jet device. **(B)** The time course of fiber deflection was well described by a single exponential (red line) with a time constant of 155 μs, corresponding to a rise time (5–95%) of 465 μs. This time constant was set by the relaxation time $\tau_F = \lambda_F/k_F$ = 160 μs of the fiber, in which $k_F$ = 1.1 mN/m and $\lambda_F$ = 173 nN·s/m were the stiffness and friction coefficient of the fiber, respectively. **(C)** Force on the fiber as a function of the command voltage to the fluid jet. The slope of the linear relation (linear fit in red) provided the calibration constant $C$, here $C = 42$ pN/V. Same data as in **A** and **B**.

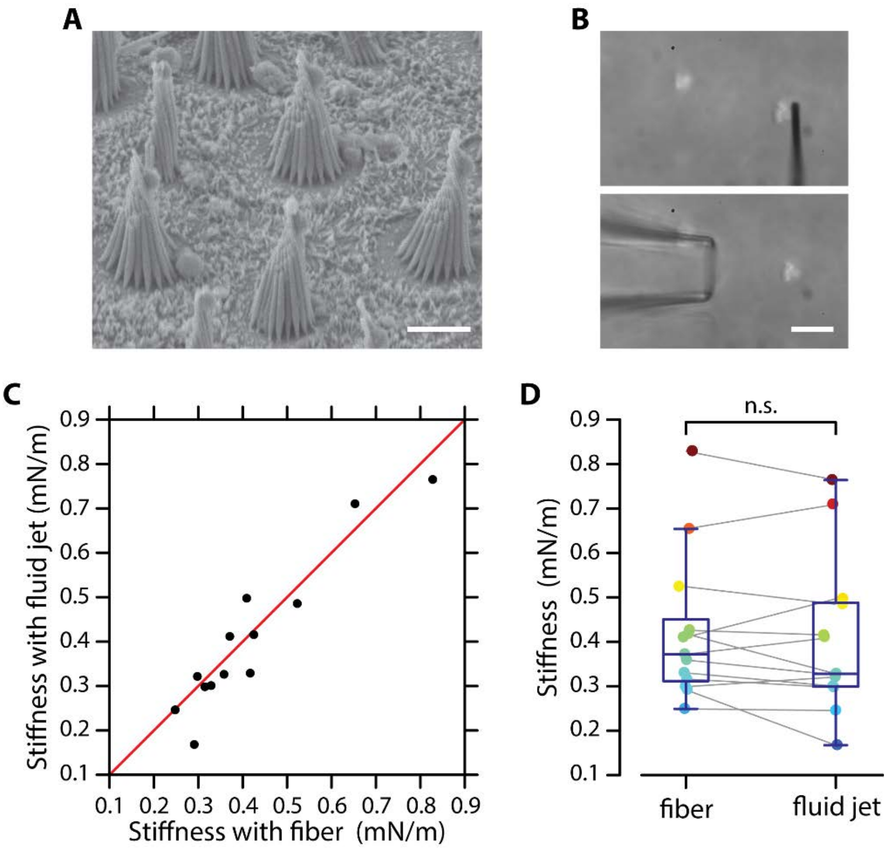


**Figure 2–figure supplement 4: Test of fluid-jet calibration in the frog's sacculus.** **(A)** Electron micrograph of hair bundles from a sacculus of the frog (strain 'Rivan92'of *Rana ridibunda*). **(B)** Micrograph of a hair bundle ready to be stimulated by a flexible fiber (top) or by a fluid jet (bottom). **(C)** The stiffness estimated from fluid-jet stimulation is here plotted as a function of that estimated from fiber stimulation for a sample of 13 hair bundles. The red line has a slope unity. **(D)** The same data as that shown in **C** is represented here as box plots on top of the data points for each method of mechanical stimulation. Grey lines connect stiffness estimates for the same hair bundles. The mean values of both distributions, 0.40±0.17 mN/m (mean±SD; median: 0.33 mN/m) and 0.42±0.16 mN/m (mean±SD; median: 0.37 mN/m) for fluid-jet stimulation and fiber stimulation, respectively, are not statistically different (paired-sample t-test; ***$p<0.001$). Scale bars in **A-B:** 5 µm.

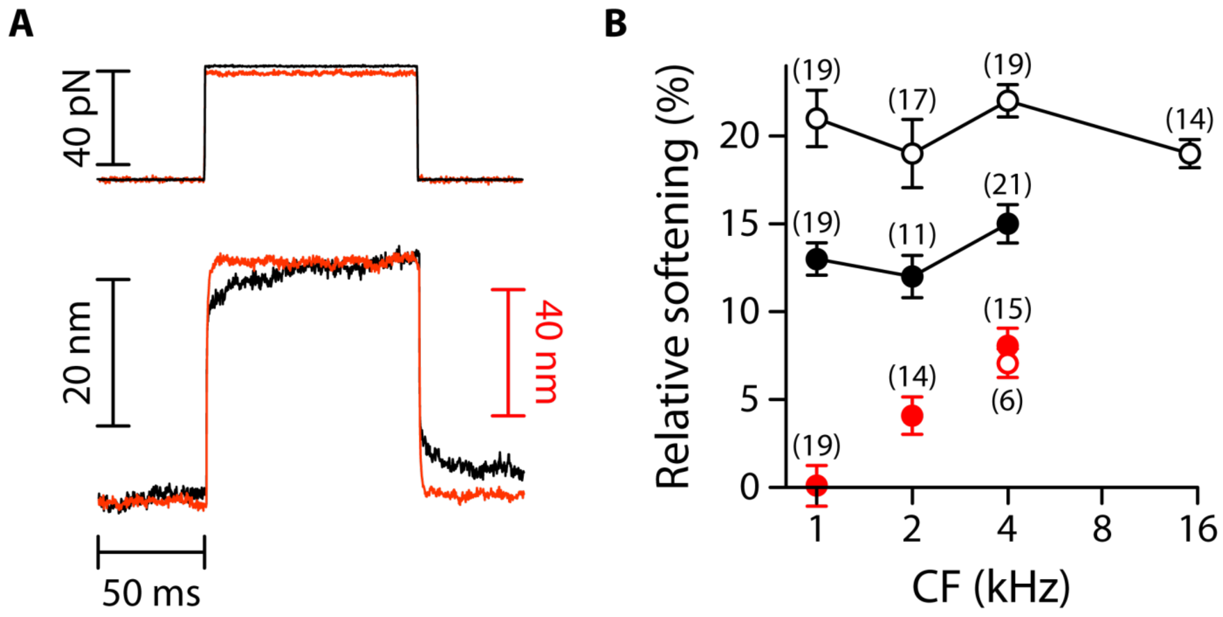


**Figure 2–figure supplement 5: Mechanical creep during a force step.**

**(A)** Hair-bundle displacement (bottom) in response to a force step (top) before (black) and after (red) tip-link disruption, here for two different outer hair cells at the same cochlear location. With intact tip links (black), the hair bundles showed a mechanical creep corresponding to a motion in the direction of the applied force over the duration of the stimulus. Remarkably, the mechanical creep was abolished by tip-link disruption (red), suggesting that the creep was associated with mechanical adaptation (Hudspeth and Gillespie, 1994). **(B)** Relative softening $(K_1 - K_2)/K_1$ associated with the mechanical creep for inner (circles) and outer (disks) hair cells before (black) and after (red) tip-link disruption. Here, $K_1$ and $K_2$ correspond to the hair-bundle stiffness measured 5–10 ms and 70–90 ms after the onset of the force step, respectively. To disrupt the tip links, the hair cells were here immersed for 15 min in a standard saline supplemented with 5-mM EDTA. The mechanical responses were then measured in standard saline. Error bars correspond to mean values ± SEM, with the number of cells indicated on the figure.

| | ANOVA | | IHC | | | | | |
|---|---|---|---|---|---|---|---|---|
| | IHC | OHC | 1-2 kHz | 1-4 kHz | 1-15 kHz | 2-4kHz | 2-15 kHz | 4-15 kHz |
| $K_{HB}$ | ********p* **=** $\mathbf{5.0\times10^{-13}}$ | ********p* **=** $\mathbf{1.7\times10^{-15}}$ | *p* = $3.8\times10^{-1}$ | ********p* **=** $\mathbf{6.4\times10^{-5}}$ | ********p* **=** $\mathbf{3.1\times10^{-8}}$ | ********p* **=** $\mathbf{1.5\times10^{-4}}$ | ********p* **=** $\mathbf{1.8\times10^{-7}}$ | ****** *p* **=** $\mathbf{5.1\times10^{-3}}$ |

| | OHC | | | OHC/IHC | | | Gradient OHC *vs.* gradient IHC |
|---|---|---|---|---|---|---|---|
| | 1-2 kHz | 1-4 kHz | 2-4 kHz | 1 kHz | 2 kHz | 4 kHz | |
| $K_{HB}$ | ****** *p* **=** $\mathbf{1.7\times10^{-3}}$ | ********p* **=** $\mathbf{9.2\times10^{-12}}$ | ********p* **=** $\mathbf{2.7\times10^{-9}}$ | ****** *p* **=** $\mathbf{7.4\times10^{-3}}$ | ********p* **=** $\mathbf{4.4\times10^{-5}}$ | ********p* **=** $\mathbf{4.5\times10^{-9}}$ | ***** *p* **=** $\mathbf{1.8\times10^{-2}}$ |

**Figure 2−table supplement 1: Statistical significance.**

The table lists p-values resulting, respectively, from a one-way ANOVA to assay statistical significance of the measured mean-value variation of the hair-bundle stiffness $K_{HB}$ between different cochlear locations for inner (IHC) and outer (OHC) hair cells, from two-tailed unpaired Student's *t*-tests with Welch’s correction to compare mean values of $K_{HB}$ between two groups of a given hair-cell type (IHC or OHC) with different characteristic frequencies (CF) or between the two cell types (OHC/IHC) when they are associated to the same characteristic frequency. The last entry (Gradient OHC *vs.* gradient IHC) provides the p-value to assay the statistical significance between the slopes of a weighted linear regression of the relation between $K_{HB}$ and the characteristic frequency of the hair cell. A bold font was used to help find statistically significant differences.

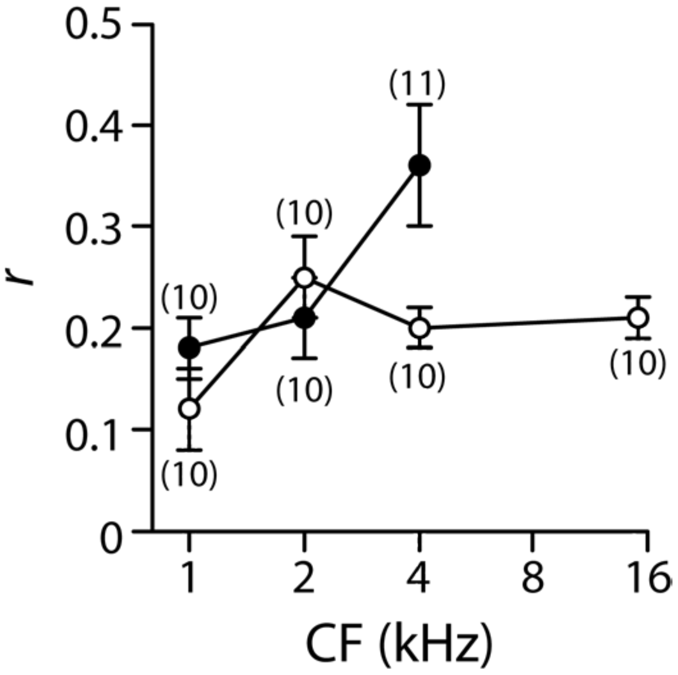


**Figure 3–figure supplement 1: Gating-spring contribution to the hair-bundle stiffness.**

We measured the amplitude of the hair-bundle movement evoked by a sinusoidal fluid-jet stimulus before (amplitude $X_1$) and after (amplitude $X_2$) application of EDTA iontophoresis (see an example in Fig 4A of the main text). The ratio $r = (X_2 - X_1)/X_2$ is here plotted as a function of characteristic frequency (CF) for inner (white disks) and outer (black disks) hair cells. Because EDTA disrupted the tip links, the ratio $r = \mathrm{K_{GS}}/\mathrm{K_{HB}}$ quantifies the contribution $\mathrm{K_{GS}}$ of the gating springs to the stiffness $\mathrm{K_{HB}}$ of an intact bundle; the complementary ratio, $1 - r = \mathrm{K_{SP}}/\mathrm{K_{HB}}$, represents the relative contribution of the stereociliary pivots, of stiffness $\mathrm{K_{SP}}$. Error bars correspond to mean values ± SEM, with the number of cells indicated on the figure. These experiments were performed in low-$Ca^{2+}$ saline.

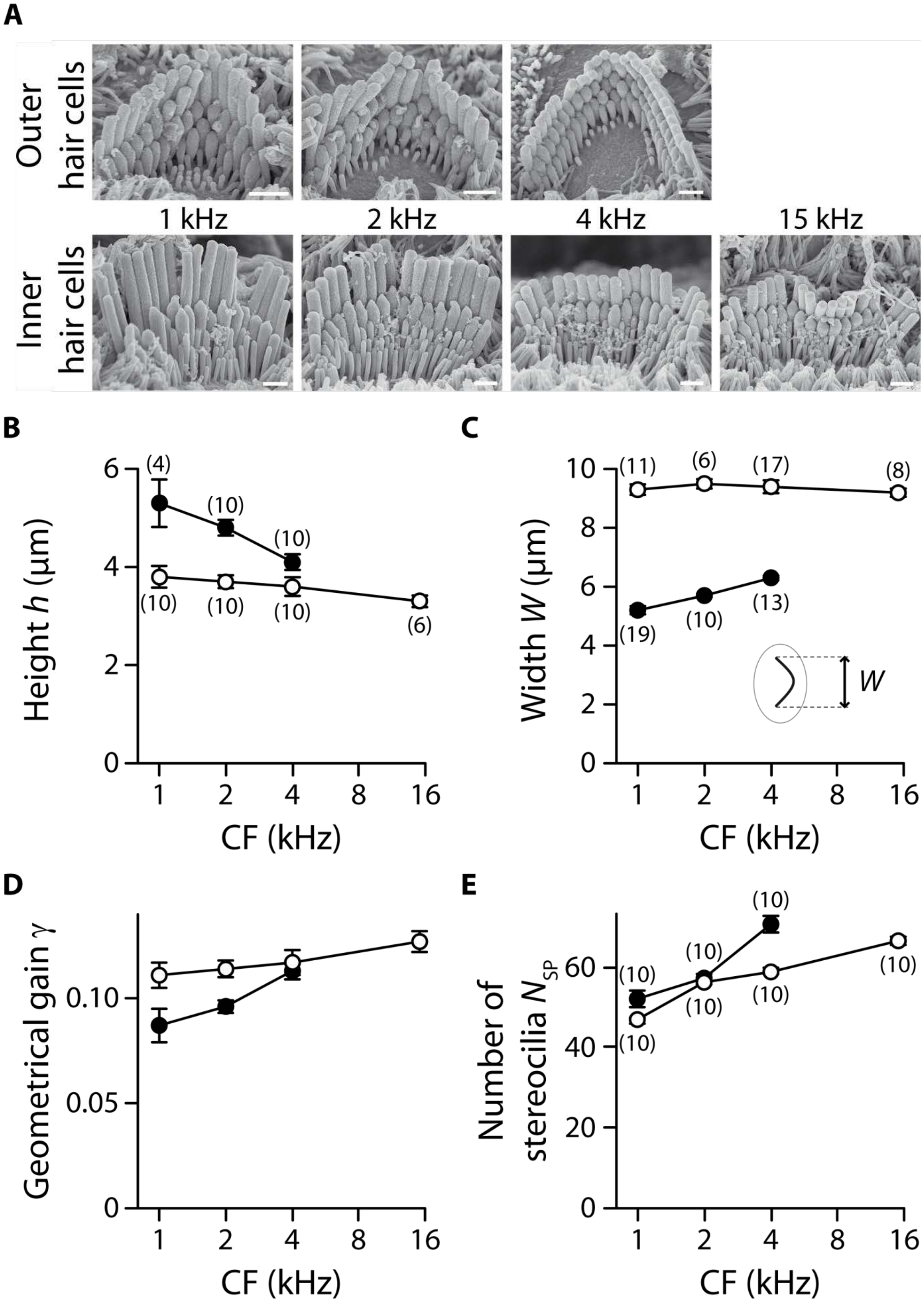


**Figure 3–figure supplement 2: Hair-bundle morphology along the tonotopic axis.**

We used electron microscopy to count the number of stereocilia in a hair bundle. To avoid dehydration artefacts, we used instead optical microscopy to measure the width and height of the hair bundle. To measure width, the hair bundle was visualized from above (Fig. 1C). To measure height, the hair bundle was visualized from the side after it had been pushed against the apical surface of the hair cell with a glass rod. **(A)** Electron micrographs of typical outer (top row) and inner (bottom row) hair cells at cochlear locations corresponding to characteristic frequencies indicated on the figure. Tension in the tip links results in stereociliary tips with a prolate shape, which is also indicative of tip-link orientation. Scale bars: 1 µm. The height $h$ **(B)**, width $W$ **(C)**, geometrical gain $\gamma$ **(D)**, and number of stereocilia $N_{SP}$ **(E)** of a hair bundle are plotted as a function of the characteristic frequency (CF) for inner (white disks) and outer (black disks) hair cells. In panel **C**, the inset shows a schematic top view of a hair bundle and indicates how the bundle's width was measured. The geometrical gain was calculated as the ratio of the interstereociliary spacing and the hair-bundle height, assuming an interstereociliary spacing of 462 nm for all outer hair cells and 420 nm for all inner hair cells. The error bars in **B**, **C** and **E** represent standard errors of the means with numbers of cells indicated between brackets; in **D**, error bars are calculated as described in the Methods.

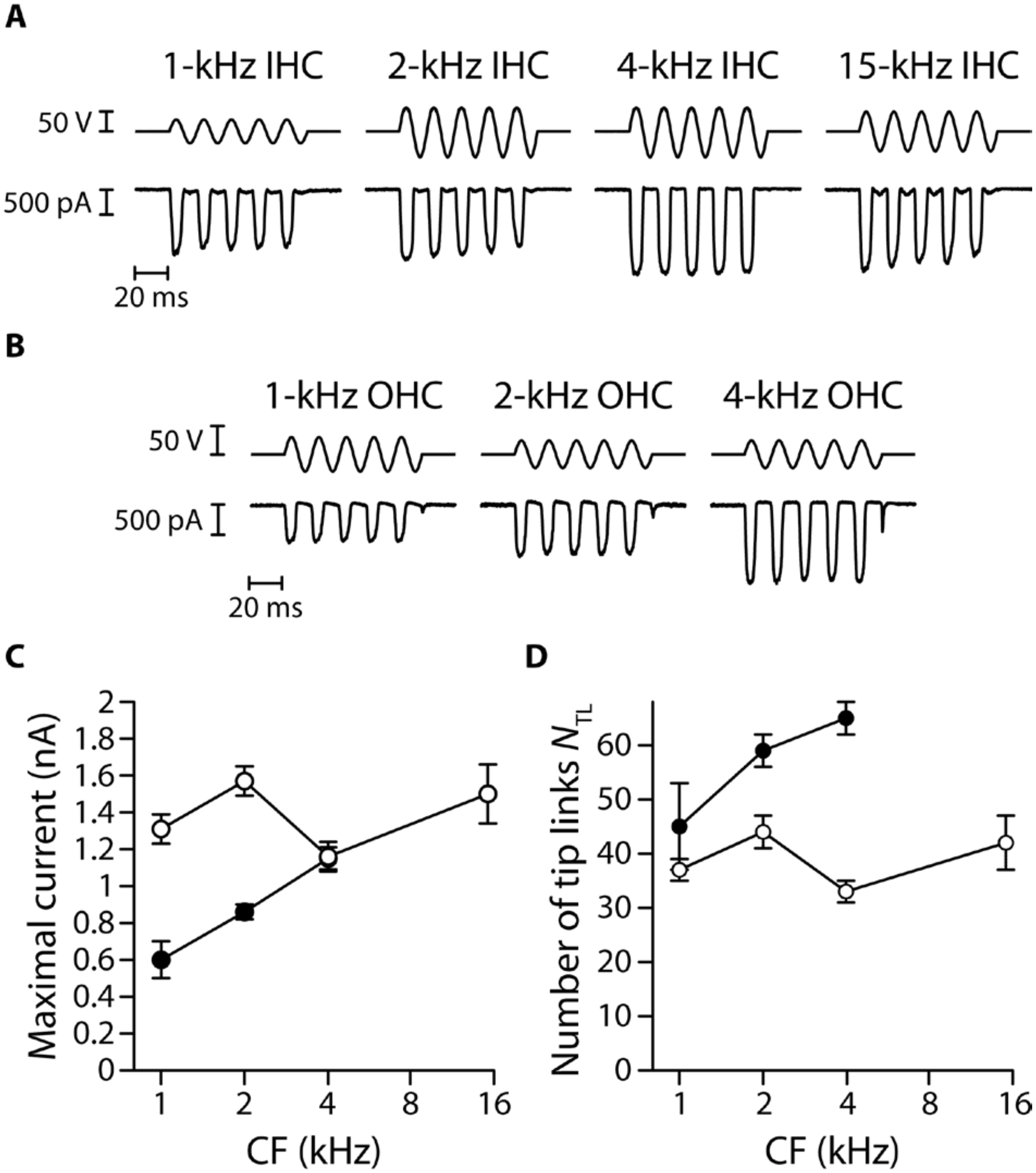


**Figure 3–figure supplement 3: Transduction currents and number of intact tip links along the tonotopic axis.**

A large mechanical stimulus (top; fluid-jet voltage command) was applied to the hair bundle to estimate the magnitude of the transduction current (bottom) at saturation in inner hair cells **(A)** and outer hair cells **(B)** along the tonotopic axis of the cochlea. Maximal current **(C)** and estimated number $N_{\mathrm{TL}}$ of tip links (**(D)**; see Methods) are plotted as a function of the characteristic frequency (CF) for inner (white disks) and outer (black disks) hair cells. Error bars correspond to mean values ± SEM, with n = 10 cells for all data points. In panels **A** and

**B**, the hair cell's characteristic frequency is indicated above each recording; the frequency of stimulation was 60 Hz. In these experiments, the hair cells were immersed in standard saline at a $Ca^{2+}$ concentration of 1.5 mM.

| | Characteristic frequency (kHz) | 1 | 2 | 4 | 15 |
|---|---|---|---|---|---|
| Inner hair cells | Width $W$ (µm) | 9.3 ± 0.2 (n=11) | 9.5 ± 0.2 (n=6) | 9.4 ± 0.2 (n=17) | 9.2 ± 0.1 (n=8) |
| | Height $h$ (µm) | 3.8 ± 0.2 (n=10) | 3.7 ± 0.1 (n=10) | 3.6 ± 0.2 (n=10) | 3.3 ± 0.1 (n=6) |
| | Number of stereocilia $N_{SP}$ | 46.6 ± 1.1 (n=10) | 56.4 ± 0.6 (n=10) | 59.0 ± 0.8 (n=10) | 66.6 ± 0.9 (n=10) |
| | Effective radius $R_{HB}$ (µm) | 2.9 | 2.9 | 2.9 | 2.7 |
| Outer hair cells | Width $W$ (µm) | 5.2 ± 0.1 (n=19) | 5.7 ± 0.1 (n=10) | 6.3 ± 0.1 (n=13) | N.A. |
| | Height $h$ (µm) | 5.3 ± 0.5 (n=4) | 4.8 ± 0.2 (n=10) | 4.1 ± 0.2 (n=10) | N.A. |
| | Number of stereocilia $N_{SP}$ | 52.0 ± 2.0 (n=10) | 57.0 ± 1.3 (n=10) | 70.6 ± 1.7 (n=10) | N.A. |
| | Effective radius $R_{HB}$ (µm) | 2.6 | 2.6 | 2.5 | N.A |

**Figure 3−table supplement 1: Morphological parameters of inner and outer hair-cell bundles.**

Data correspond to mean values ± SEM, with the number of cells indicated in parentheses. The width $W$ and height $h$ of the hair bundle are used to calculate an effective hydrodynamic radius $R_{HB}$ (see Eq. 1 in Methods) and plotted as a function of the hair cell's characteristic frequency in Figure 3−figure supplement 2.

| | ANOVA | | IHC | | | | | |
|---|---|---|---|---|---|---|---|---|
| | IHC | OHC | 1-2 kHz | 1-4 kHz | 1-15 kHz | 2-4kHz | 2-15 kHz | 4-15 kHz |
| $r$ | $p = 7.6\times10^{-2}$ | *$\boldsymbol{p} = \mathbf{2.0\times10^{-2}}$ | $p = 5.3\times10^{-2}$ | $p = 1.2\times10^{-1}$ | $p = 9.2\times10^{-2}$ | $p = 3.2\times10^{-1}$ | $p = 4.3\times10^{-1}$ | $p = 7.4\times10^{-1}$ |
| $K_{GS}$ | ***$\boldsymbol{p} = \mathbf{1.3\times10^{-5}}$ | ***$\boldsymbol{p} = \mathbf{1.7\times10^{-6}}$ | *$\boldsymbol{p} = \mathbf{2.1\times10^{-2}}$ | ***$\boldsymbol{p} = \mathbf{8.9\times10^{-5}}$ | ***$\boldsymbol{p} = \mathbf{3.3\times10^{-6}}$ | *$\boldsymbol{p} = \mathbf{4.2\times10^{-2}}$ | ***$\boldsymbol{p} = \mathbf{4.4\times10^{-4}}$ | *$\boldsymbol{p} = \mathbf{3.9\times10^{-2}}$ |
| $K_{SP}$ | ***$\boldsymbol{p} = \mathbf{2.0\times10^{-14}}$ | ***$\boldsymbol{p} = \mathbf{8.3\times10^{-8}}$ | $p = 6.6\times10^{-1}$ | ***$\boldsymbol{p} = \mathbf{1.9\times10^{-4}}$ | ***$\boldsymbol{p} = \mathbf{1.5\times10^{-8}}$ | ***$\boldsymbol{p} = \mathbf{3.6\times10^{-5}}$ | ***$\boldsymbol{p} = \mathbf{1.3\times10^{-8}}$ | **$\boldsymbol{p} = \mathbf{4.1\times10^{-3}}$ |
| $\kappa$ | **$\boldsymbol{p} = \mathbf{1.0\times10^{-3}}$ | $p = 7.7\times10^{-1}$ | $p = 1.6\times10^{-1}$ | $p = 7.0\times10^{-2}$ | *$\boldsymbol{p} = \mathbf{1.4\times10^{-2}}$ | **$\boldsymbol{p} = \mathbf{1.6\times10^{-3}}$ | ***$\boldsymbol{p} = \mathbf{3.9\times10^{-5}}$ | $p = 7.3\times10^{-1}$ |
| $k_{GS}$ | *$\boldsymbol{p} = \mathbf{3.0\times10^{-2}}$ | **$\boldsymbol{p} = \mathbf{1.9\times10^{-3}}$ | $p = 3.0\times10^{-1}$ | **$\boldsymbol{p} = \mathbf{1.6\times10^{-3}}$ | **$\boldsymbol{p} = \mathbf{1.8\times10^{-3}}$ | *$\boldsymbol{p} = \mathbf{1.5\times10^{-2}}$ | *$\boldsymbol{p} = \mathbf{1.6\times10^{-2}}$ | $p = 1$ |

| | OHC | | | OHC/IHC | | | Gradient OHC *vs.* gradient IHC |
|---|---|---|---|---|---|---|---|
| | 1-2 kHz | 1-4 kHz | 2-4 kHz | 1 kHz | 2 kHz | 4 kHz | |
| $r$ | $p = 5.8\times10^{-1}$ | *$\boldsymbol{p} = \mathbf{1.3\times10^{-2}}$ | $p = 5.5\times10^{-2}$ | $p = 2.7\times10^{-1}$ | $p = 5.3\times10^{-1}$ | *$\boldsymbol{p} = \mathbf{2.1\times10^{-2}}$ | *$\boldsymbol{p} = \mathbf{2.8\times10^{-2}}$ |
| $K_{GS}$ | $p = 1.5\times10^{-1}$ | ***$\boldsymbol{p} = \mathbf{2.2\times10^{-4}}$ | ***$\boldsymbol{p} = \mathbf{7.1\times10^{-4}}$ | *$\boldsymbol{p} = \mathbf{1.4\times10^{-2}}$ | $p = 1.5\times10^{-1}$ | ***$\boldsymbol{p} = \mathbf{7.0\times10^{-4}}$ | $p = 1.6\times10^{-1}$ |
| $K_{SP}$ | *$\boldsymbol{p} = \mathbf{1.8\times10^{-2}}$ | ***$\boldsymbol{p} = \mathbf{2.1\times10^{-5}}$ | ***$\boldsymbol{p} = \mathbf{9.5\times10^{-4}}$ | *$\boldsymbol{p} = \mathbf{3.9\times10^{-2}}$ | ***$\boldsymbol{p} = \mathbf{5.2\times10^{-5}}$ | ***$\boldsymbol{p} = \mathbf{9.1\times10^{-4}}$ | **$\boldsymbol{p} = \mathbf{7.6\times10^{-3}}$ |
| $\kappa$ | $p = 6.6\times10^{-1}$ | $p = 4.8\times10^{-1}$ | $p = 6.6\times10^{-1}$ | **$\boldsymbol{p} = \mathbf{8.1\times10^{-3}}$ | ***$\boldsymbol{p} = \mathbf{1.4\times10^{-9}}$ | **$\boldsymbol{p} = \mathbf{6.1\times10^{-3}}$ | $p = 1.9\times10^{-1}$ |
| $k_{GS}$ | $p = 6.9\times10^{-1}$ | **$\boldsymbol{p} = \mathbf{6.1\times10^{-3}}$ | **$\boldsymbol{p} = \mathbf{8.5\times10^{-3}}$ | $p = 8.1\times10^{-2}$ | $p = 6.3\times10^{-2}$ | *$\boldsymbol{p} = \mathbf{1.5\times10^{-2}}$ | $p = 1.5\times10^{-1}$ |

**Figure 3–table supplement 2: Statistical significance.**

The table lists p-values resulting, respectively, from a one-way ANOVA to assay statistical significance of the measured mean-value variation of a given variable between different cochlear locations for inner (IHC) and outer (OHC) hair cells, from two-tailed unpaired Student's *t*-tests with Welch's correction to compare mean values of the variable between two groups of a given hair-cell type (IHC or OHC) with different characteristic frequencies (CF) or between the two cell types (OHC/IHC) when they are associated to the same characteristic frequency. The last entry provides the p-value to assay the statistical significance between the slopes of a weighted linear regression of the relation between the variable and the characteristic frequency of the hair cell. A bold font was used to help find statistically significant differences. The variables in the table correspond to the relative contribution $r$ and the absolute contribution $K_{GS}$ of the gating springs to the hair-bundle stiffness, the contribution $K_{SP}$ of the stereociliary pivots to the hair-bundle stiffness, the rotational stiffness $\kappa$ of a single stereocilium, and the stiffness $k_{GS}$ of a single gating spring.

| | ANOVA | | IHC | | | | | |
|---|---|---|---|---|---|---|---|---|
| | IHC | OHC | 1-2 kHz | 1-4 kHz | 1-15 kHz | 2-4kHz | 2-15 kHz | 4-15 kHz |
| $\Delta X_{\mathrm{R}}$ | $p = 4.3\times10^{-1}$ | ***$\boldsymbol{p = 7.3\times10^{-4}}$ | $p = 8.5\times10^{-2}$ | $p = 4.3\times10^{-1}$ | $p = 1.8\times10^{-1}$ | $p = 5.0\times10^{-1}$ | $p = 8.7\times10^{-1}$ | $p = 6.5\times10^{-1}$ |
| $T_{\mathrm{R}}$ | ***$\boldsymbol{p = 2.6\times10^{-5}}$ | ***$\boldsymbol{p = 1.0\times10^{-4}}$ | $p = 2.1\times10^{-1}$ | *$\boldsymbol{p = 4.0\times10^{-2}}$ | **$\boldsymbol{p = 5.6\times10^{-3}}$ | $p = 1.1\times10^{-1}$ | *$\boldsymbol{p = 1.3\times10^{-2}}$ | $p = 1.6\times10^{-1}$ |
| $t_{\mathrm{R}}$ | ***$\boldsymbol{p = 6.1\times10^{-4}}$ | ***$\boldsymbol{p = 1.6\times10^{-4}}$ | $p = 6.3\times10^{-1}$ | *$\boldsymbol{p = 3.7\times10^{-2}}$ | *$\boldsymbol{p = 1.3\times10^{-2}}$ | $p = 5.5\times10^{-2}$ | *$\boldsymbol{p = 1.9\times10^{-2}}$ | $p = 6.4\times10^{-1}$ |

| | OHC | | | OHC/IHC | | | Gradient OHC *vs.* gradient IHC |
|---|---|---|---|---|---|---|---|
| | 1-2 kHz | 1-4 kHz | 2-4 kHz | 1 kHz | 2 kHz | 4 kHz | |
| $\Delta X_{\mathrm{R}}$ | *$\boldsymbol{p = 1.6\times10^{-2}}$ | **$\boldsymbol{p = 1.2\times10^{-3}}$ | *$\boldsymbol{p = 3.9\times10^{-2}}$ | $p = 1.1\times10^{-1}$ | $p = 7.1\times10^{-1}$ | *$\boldsymbol{p = 3.2\times10^{-2}}$ | ***$\boldsymbol{p = 4.9\times10^{-4}}$ |
| $T_{\mathrm{R}}$ | **$\boldsymbol{p = 3.2\times10^{-3}}$ | **$\boldsymbol{p = 1.3\times10^{-3}}$ | **$\boldsymbol{p = 7.5\times10^{-3}}$ | $p = 4.9\times10^{-1}$ | *$\boldsymbol{p = 3.5\times10^{-2}}$ | **$\boldsymbol{p = 6.6\times10^{-3}}$ | *$\boldsymbol{p = 3.7\times10^{-2}}$ |
| $t_{\mathrm{R}}$ | *$\boldsymbol{p = 3.1\times10^{-2}}$ | **$\boldsymbol{p = 2.6\times10^{-3}}$ | *$\boldsymbol{p = 1.7\times10^{-2}}$ | $p = 6.2\times10^{-1}$ | $p = 8.2\times10^{-2}$ | $p = 6.5\times10^{-2}$ | *$\boldsymbol{p = 2.0\times10^{-2}}$ |

**Figure 5–table supplement 1: Statistical significance.**

The table lists p-values resulting, respectively, from a one-way ANOVA to assay statistical significance of the measured mean-value variation of a given variable between different cochlear locations for inner (IHC) and outer (OHC) hair cells, from two-tailed unpaired Student's *t*-tests with Welch's correction to compare mean values of the variable between two groups of a given hair-cell type (IHC or OHC) with different characteristic frequencies (CF) or between the two cell types (OHC/IHC) when they are associated to the same characteristic frequency. The last entry provides the p-value to assay the statistical significance between the slopes of a weighted linear regression of the relation between the variable and the characteristic frequency of the hair cell. A bold font was used to help find statistically significant differences. The variables in the table correspond to the net positive movement $\Delta X_{\mathrm{R}}$ of the hair bundle evoked at steady state by tip-link disruption, the mechanical tension $T_{\mathrm{R}}$ in the hair bundle, and the mechanical tension $t_{\mathrm{R}}$ in a single gating spring.

| | ANOVA | | IHC | | | | | |
|---|---|---|---|---|---|---|---|---|
| | IHC | OHC | 1-2 kHz | 1-4 kHz | 1-15 kHz | 2-4kHz | 2-15 kHz | 4-15 kHz |
| $\Delta X_{\mathrm{Ca}}$ | $p = 1.9\times10^{-1}$ | $p = 4.6\times10^{-1}$ | $p = 6.8\times10^{-1}$ | $p = 9.5\times10^{-2}$ | $p = 4.1\times10^{-1}$ | *$p = 2.2\times10^{-2}$ | $p = 2.3\times10^{-1}$ | $p = 6.0\times10^{-1}$ |
| $\Delta T$ | ***$p = 6.5\times10^{-6}$ | **$p = 5.1\times10^{-3}$ | $p = 6.4\times10^{-1}$ | ***$p = 6.3\times10^{-4}$ | **$p = 5.9\times10^{-3}$ | ***$p = 2.8\times10^{-4}$ | **$p = 4.2\times10^{-3}$ | $p = 4.0\times10^{-1}$ |
| $t_{\mathrm{max}}$ | ***$p = 3.1\times10^{-6}$ | ***$p = 7.2\times10^{-4}$ | $p = 5.8\times10^{-1}$ | **$p = 1.4\times10^{-3}$ | **$p = 1.8\times10^{-3}$ | ***$p = 5.1\times10^{-4}$ | ***$p = 7.7\times10^{-4}$ | $p = 1$ |

| | OHC | | | OHC/IHC | | | Gradient OHC *vs.* gradient IHC |
|---|---|---|---|---|---|---|---|
| | 1-2 kHz | 1-4 kHz | 2-4 kHz | 1 kHz | 2 kHz | 4 kHz | |
| $\Delta X_{\mathrm{Ca}}$ | $p = 4.3\times10^{-1}$ | $p = 2.7\times10^{-1}$ | $p = 5.9\times10^{-1}$ | $p = 3.5\times10^{-1}$ | $p = 8.2\times10^{-1}$ | $p = 4.6\times10^{-1}$ | $p = 7.1\times10^{-2}$ |
| $\Delta T$ | *$p = 4.9\times10^{-2}$ | *$p = 1.0\times10^{-2}$ | $p = 5.8\times10^{-2}$ | $p = 8.8\times10^{-1}$ | **$p = 5.5\times10^{-3}$ | $p = 2.7\times10^{-1}$ | *$p = 1.7\times10^{-2}$ |
| $t_{\mathrm{max}}$ | $p = 1.3\times10^{-1}$ | **$p = 3.9\times10^{-3}$ | *$p = 2.8\times10^{-2}$ | $p = 8.4\times10^{-1}$ | *$p = 4.9\times10^{-2}$ | $p = 4.9\times10^{-1}$ | *$p = 2.2\times10^{-2}$ |

**Figure 6–table supplement 1: Statistical significance.**

The table lists p-values resulting, respectively, from a one-way ANOVA to assay statistical significance of the measured mean-value variation of a given variable between different cochlear locations for inner (IHC) and outer (OHC) hair cells, from two-tailed unpaired Student's *t*-tests with Welch's correction to compare mean values of the variable between two groups of a given hair-cell type (IHC or OHC) with different characteristic frequencies (CF) or between the two cell types (OHC/IHC) when they are associated to the same characteristic frequency. The last entry provides the p-value to assay the statistical significance between the slopes of a weighted linear regression of the relation between the variable and the characteristic frequency of the hair cell. A bold font was used to help find statistically significant differences. The variables in the table correspond to the negative hair-bundle movement $\Delta X_{\mathrm{Ca}}$, the corresponding increase in hair-bundle tension $\Delta T$, and the maximal tension $t_{\mathrm{max}}$ in a single gating spring evoked by EDTA iontophoresis just before tip-link disruption.
